# Mechanical control of innate immune responses against viral infection revealed in a human Lung Alveolus Chip

**DOI:** 10.1101/2021.04.26.441498

**Authors:** Haiqing Bai, Longlong Si, Amanda Jiang, Chaitra Belgur, Roberto Plebani, Crystal Oh, Melissa Rodas, Atiq Nurani, Sarah Gilpin, Rani K. Powers, Girija Goyal, Rachelle Prantil- Baun, Donald E. Ingber

**Author notes:** These authors contribute equally to this work (Haiqing Bai, Longlong Si).

## Abstract

Mechanical forces associated with breathing play a fundamental role in lung development and disease but the molecular pathways remain largely unknown. Here, we used a mechanically actuatable Human Lung Alveolus Chip that recapitulates human lung alveolar type I and type II cell differentiation, alveolar-capillary interface formation, and genome-wide gene expression profiles characteristic of the distal lung to investigate the role of physical forces associated with cyclic breathing motions in lung innate immune responses to viral infection. When the mechanically active Alveolus Chips are infected with the influenza H3N2 virus, a cascade of host responses is elicited on-chip, including increased production of cytokines and expression of inflammation-associated genes in pulmonary epithelial and endothelial cells, resulting in enhanced recruitment of circulating immune cells as occurs during viral infection in vivo. Surprisingly, studies carried out in parallel with static chips revealed that physiological breathing motions suppress viral replication by activating protective innate immune responses in epithelial and endothelial cells. This is mediated at least in part through upregulation of S100 calcium-binding protein A7 (S100A7), which binds to the Receptor for Advanced Glycation End Products (RAGE), an inflammatory mediator that is most highly expressed in the lung alveolus *in vivo*. This mechano-immunological control mechanism is further supported by the finding that existing RAGE inhibitor drugs can suppress the production of inflammatory cytokines in response to influenza virus infection in this model. S100A7-RAGE interactions and modulation of mechanical ventilation parameters could therefore serve as new targets for therapeutic intervention in patients infected with influenza and other potential pandemic viruses that cause life-threatening lung inflammation.

## INTRODUCTION

Innate immune responses in the lung serve as a first line of defense against infections by respiratory viruses, such as influenza A viruses and SARS-CoV-2, yet we know little about how these responses are controlled locally within the physical microenvironment of the human breathing lung. One reason is that it is difficult, if not impossible, to tease out the regulatory cues in the local environment within a living organ in vivo, and the mechanical cues experienced human lung are absent in conventional culture models.

Breathing is central to life, but its biological effects in lung beyond oxygenation are poorly understood. Breathing motions in human lung exert dynamic physical forces on its constituent cells and tissues that promote lung branching^1^ and alveolar differentiation^2^ during normal lung development, and contribute to the etiology of various lung diseases, including acute lung injury^3, 4^, pulmonary edema^5^, and pulmonary fibrosis^6^. Mechanical forces have been implicated in activation of neutrophils, monocytes, and macrophages as well^7^. However, nothing is known about how breathing motions and mechanical forces impact innate immunity and responses to infection within lung parenchymal cells and pulmonary microvascular endothelium in an organ-relevant context. This is in large part because of the lack of experimental models that can mimic human lung structures and pathophysiology at the tissue and organ levels while permitting control over breathing motions.

Here we use a human organ-on-a-chip (Organ Chip) microfluidic model of the lung alveolus (Alveolus Chip) that recapitulates the human alveolar-capillary interface with an air-liquid interface (ALI) and vascular fluid flow while allowing independent control over breathing motions^4, 5, 8^ to address this challenge *in vitro*. Human Alveolus Chips have been previously shown to faithfully mimic human lung physiology, disease states, and adenoviral vector-mediated gene therapy delivery, as well as therapeutic efficacy and toxicity responses^4, 5, 8, 9^. To explore whether breathing motions influence host innate immunity and responses to infection, we used influenza H3N2 virus as a pathogen because it is one of the predominant strains in circulation and a leading cause of lower respiratory infection and hospitalization^10^. Rapid mutation and reassortment of influenza A viruses also have a high likelihood of leading to the emergence of novel strain that can cause future pandemics, which constitutes an ongoing threat to public health.

Influenza A viruses that reach the alveolus damage the tissue barrier (alveolar-endothelial interface), induce pulmonary edema, recruit inflammatory cells, and if uncontrolled, cause severe viral pneumonia and acute respiratory distress syndrome, often leading to death^11^.

Current studies of influenza infection in the distal lung rely on human cell lines that fail to recapitulate disease pathogenesis or mouse models that do not reflect the anatomy or pathophysiology of the natural human host. Thus, use of human Alveolus Chips can enable us to address questions relating to human lung responses to viral infection and contributions of related breathing motions, which would be impossible in other preclinical experimental models.

## RESULTS

### Recapitulation of alveolar cell differentiation and alveolar-capillary interface formation

The Human Alveolus Chip used in this study is a microfluidic device containing two parallel channels separated by an extracellular matrix (ECM)-coated porous membrane lined by primary human lung alveolar epithelium cells cultured under an ALI on its upper surface and primary human pulmonary microvascular endothelial cells on the lower surface, which are fed by continuous flow of culture medium through the lumen of the lower vascular channel (**Fig. 1a**)^8, 12, 13^. The engineered alveolar-capillary interface is also exposed to cyclic mechanical deformations (5% cyclic strain at 0.25Hz) to mimic physiological breathing motions at tidal volume^14^ and the normal respiratory rate of humans via application of cyclic suction to hollow side chambers within the flexible polydimethylsiloxane (PDMS) device (**Fig. 1a**).

**Figure 1.**
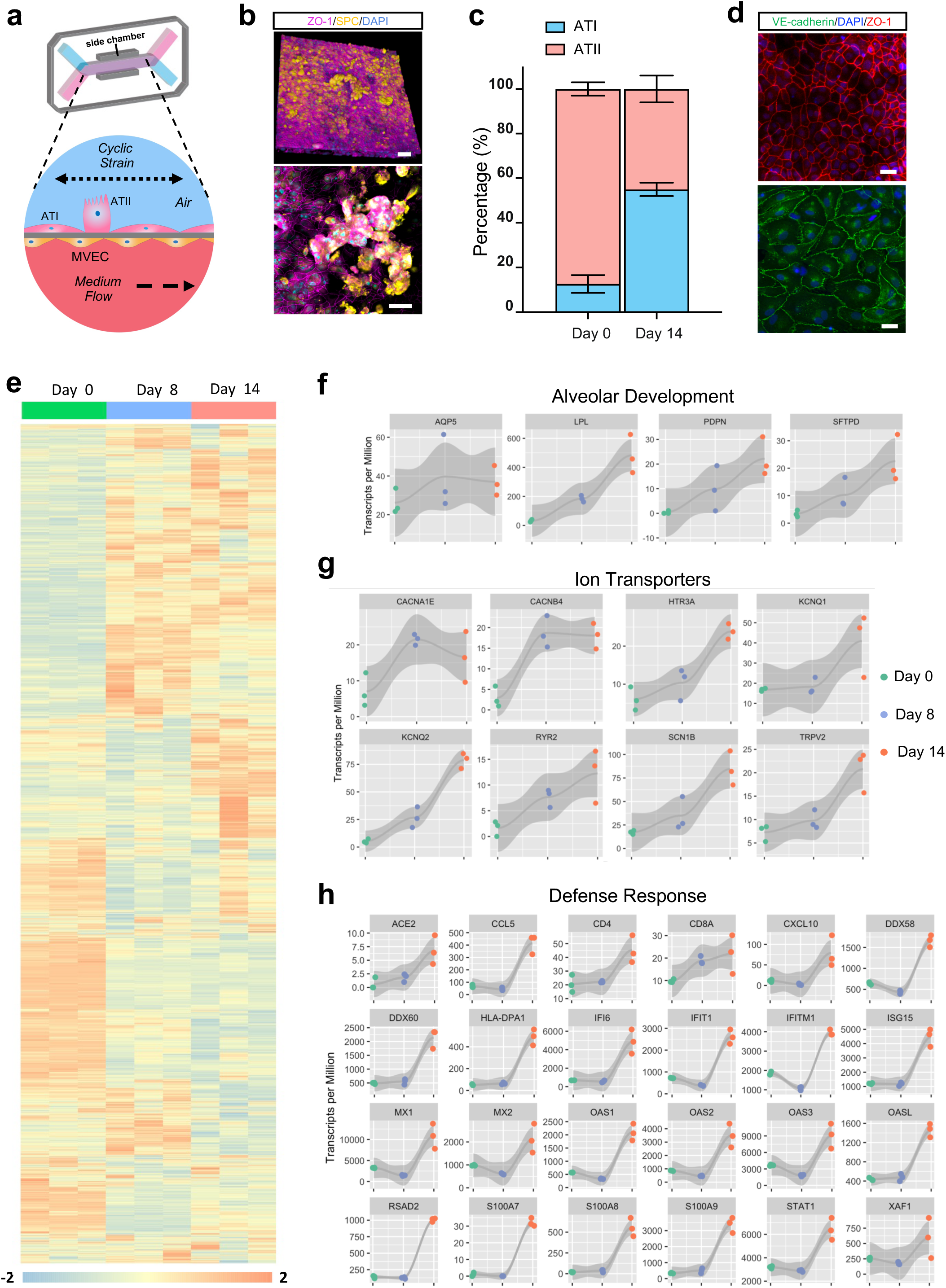
The Human Alveolus Chip. (**a**) Schematic of Human Alveolus Chip with primary alveolar epithelial type I (ATI) and type II (ATII) cells lining the upper surface of the porous ECM-coated membrane in the air channel with and pulmonary microvascular endothelial cells (MVEC) on the lower surface of the same membrane in the basal vascular channel that is continuously perfused with medium. The entire membrane and adherent alveolar-capillary interface are exposed to physiological cyclic strain by applying cyclic suction to neighboring hollow chambers (grey) within the flexible PDMS microfluidic device. (**b**) Two magnifications of immunofluorescence micrographs showing the distribution of ZO1-containing tight junctions and ATII cell marker surfactant protein C (*SPC*) in the epithelium of the Alveolus Chip (bar, 50 μm). (**c**) Graph showing the percentages of ATI and ATII cells at the time of plating and 14 days after culture on-chip. Data represent mean ± SD; n = 3 biological replicates. (**d**) Immunofluorescence micrographs showing alveolar epithelial cells (top) and endothelial cells (bottom) within the Alveolus Chip stained for ZO1 and VE-cadherin, respectively (bar, 50 μm). (**e**) Temporal gene expression profiles in the alveolar epithelial cells on-chip. (**f-h)** Time course of expression of selected genes that are involved in (**f**) alveolar development, (**g**) ion-transport and (**h**) defense response in epithelial cells cultured for up to two weeks on-chip; n = 3 biological replicates.

Using a commercial source of primary human lung alveolar epithelial cells composed of 90% alveolar type II (ATII) cells (**Supplementary Fig. 1a**) and an optimized ECM coating (200 µg/ml collagen IV and 15 µg/ml laminin), we found that the Alveolus Chips support differentiation of the ATII cells into to alveolar type I (ATI) cells (**Fig. 1b and Supplementary Fig. 1b**) resulting in a final ratio of ATII to ATI cells of 55:45 as demonstrated by immunostaining of ATII cell marker surfactant protein C (*SPC*) and the ATI cell marker *RAGE* (**Fig. 1c**). This is also accompanied by formation of a tight epithelial barrier (**Supplementary Fig. 1c**), which approximates that observed *in vivo*^15^. Confluent monolayers of interfaced alveolar epithelium and endothelium also display continuous cell-cell junctions (stained apically with *ZO-1* in epithelium and laterally with VE-cadherin, respectively) by 14 days of culture (**Fig. 1d**), which are maintained for at least 5 weeks on-chip.

RNA sequencing (RNA-seq) analysis of the alveolar epithelium showed that epithelial cells cultured on-chip exhibit extensive changes in mRNA expression over 14 days in culture (**Fig. 1e**) and display distinct transcriptome changes compared to the same cells maintained in conventional static Transwell cultures, as demonstrated by principal component analysis (PCA) and differential gene expression analysis (**Supplementary Fig. 2**).

Using short time-series expression miner (STEM)^16^, we identified five gene expression patterns that exhibited statistically significant changes over time when cultured on-chip in the presence of breathing-like motions (**Fig. 1e and Supplementary Fig. 3**). Gene ontology (GO) analysis revealed that many of the upregulated genes are involved in biological processes relevant to differentiation, including anatomical structure development, cell adhesion, movement of cell or subcellular components, and ECM organization (**Supplementary Fig. 3**). Examples include aquaporin 5 (*AQP5*) and podoplanin (*PDPN*) (**Fig. 1f**) that are known markers for ATI cells^17^, confirming the differentiation from ATII to ATI cells that we observed by immunostaining (**Fig. 1b and Supplementary Fig. S1b**). Lipoprotein lipase (*LPL*) and surfactant protein D (*SFTPD*), two genes involved in surfactant production, also increased (**Fig. 1f**), consistent with the previous finding that mechanical strain promotes enhanced surfactant secretion in vivo^18^ and on-chip^4^.

The upregulation of several ion channels and transporters (**Fig. 1g**) suggests that alveolar fluid clearance, which is essential for maintenance of air-liquid interface, may be increased as well. A large number of genes belonging to defense responses also were upregulated from day 8 to day 14 (**Fig. 1h**), including angiotensin converting enzyme 2 (*ACE2*), the receptor for SARS-CoV-2, which is also a key mediator of lung physiology and pathophysiology^19^. Importantly, this upregulation of alveolar development and maturation on-chip is similar to that seen in both human and mouse during the transition from fetus to birth^20, 21^.

Significant genetic reprogramming was also observed in endothelial cells cultured in the presence of cyclic mechanical strain on-chip over 14 days in culture (**Supplementary Fig. 4a, b**). The Wnt signaling pathway is essential for cross-talk between endothelium and epithelium during lung development^22^ and consistent with this, we found a number of Wnt ligands, including *WNT3*, *WNT7A*, *WNT7B*, *WNT9A*, and *WNT10A,* exhibit elevated expression in endothelial cells on-chip in the presence of breathing motions (**Supplementary Fig. 4c**). Together, our analysis reveals that human Lung Alveolus Chips that are exposed to physiological breathing motions recapitulate perinatal maturation of the alveolar-capillary interface, cell-ECM interactions, and epithelial-endothelial crosstalk that are indispensable for the development, homeostasis, and regeneration of the lung alveolus^23^.

### Influenza A Virus Infection of the Human Lung Alveolus Chip

Influenza A virus entry is mediated by hemagglutinin binding to sialic acid receptor on host cells, with avian and mammalian viral strains preferentially binding to α−2,3- or α−2,6- linked sialylated glycans, respectively^24^. Using glycan-specific lectins, we found that the human lung alveolar epithelial cells predominantly express α−2,3-linked sialic acid receptors on their apical surface on-chip (**Fig. 2a**). Consistent with this finding and results obtained in human lung tissues in vivo^25^, we found that influenza A/HongKong/8/68 (HK/68; H3N2) and A/HongKong/156/1997 (HK/97; H5N1) viruses successfully infect the epithelium in the human Alveolus Chip, whereas the influenza A/WSN/33 (WSN; H1N1) virus does not, as detected by immunostaining of viral nuclear protein (NP) (**Fig. 2b**). The specificity of this response is further exemplified by the finding that H1N1 virus that preferentially infects large airways also infects human Lung Airway Chips lined by bronchial epithelium much more effectively than H3N2^26^.

**Figure 2.**
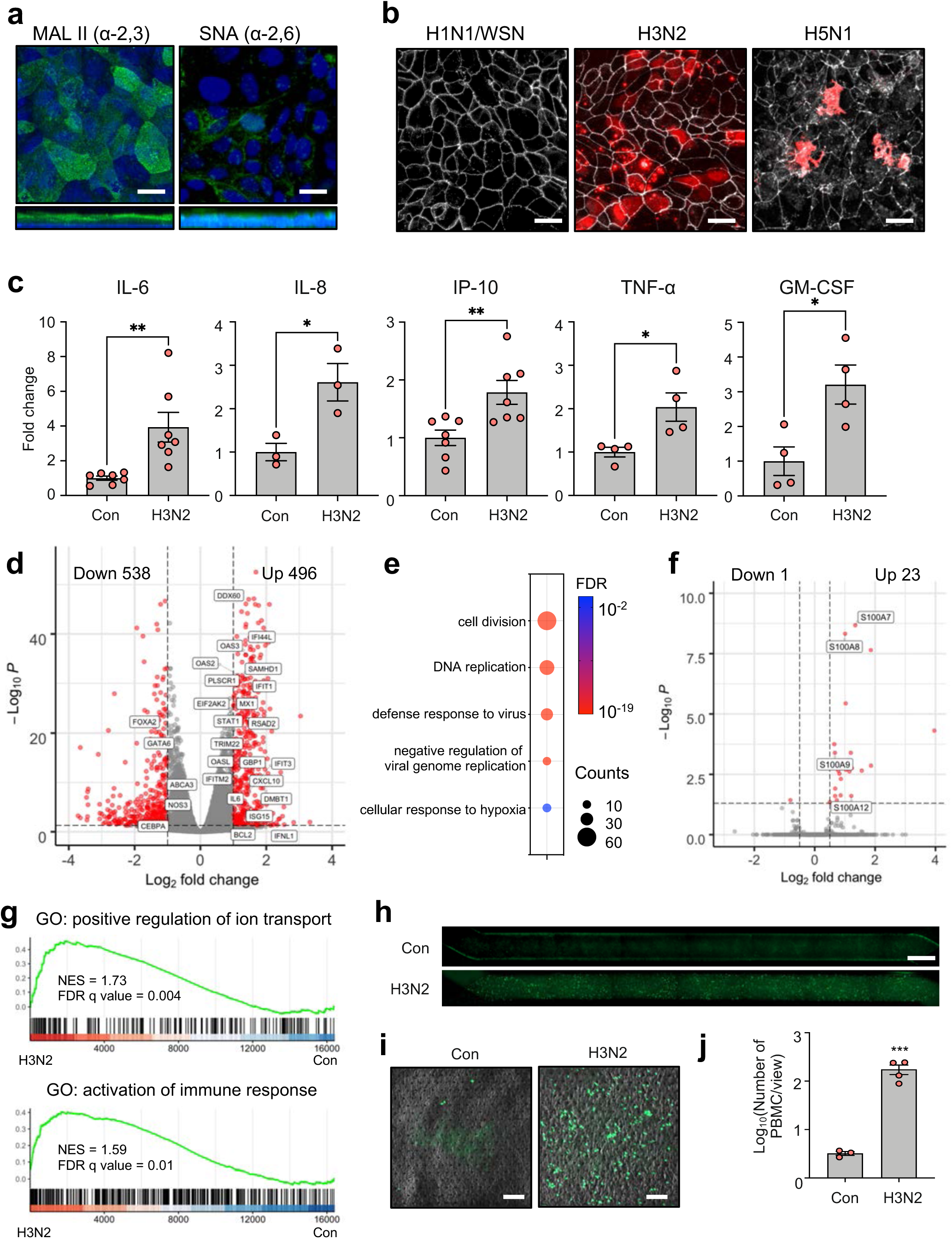
Influenza A virus infection in the Human Alveolus Chip. (**a**) Top and side fluorescence views of the alveolar epithelium cultured on-chip and stained for α-2,3-linked sialic α-2,6-linked sialic acid using Maackia Amurensis Lectin II (MAL II) and Sambucus nigra agglutinin (SNA), respectively (bar, 50 µm). (**b**) Top fluorescence views of the alveolar epithelium on-chip infected with three different influenza virus strains (WSN (H1N1), HK/68 (H3N2), or HK/97 (H5N1) at MOI = 1) and stained for viral nuclear protein (NP) (bar, 50 µm). (**c**) Levels of the indicated cytokines measured in the effluent of the vascular channels of Alveolus Chips infected with HK/68 (H3N2) virus (MOI = 1) versus control untreated chips (Con) at 48 hpi. Data represent mean ± SD; n = 3-6 biological replicates from two independent experiments; unpaired two-tailed t-test, *p<0.05, **p<0.01. (**d**) Volcano plot of DEGs in epithelial cells from HK/68 (H3N2)-infected Alveolus Chips (MOI = 1) compared to control uninfected chips (DEGs related to antiviral interferon responses are shown). (**e**) Dot plot visualization of enriched biological processes in epithelial cells of HK/68 (H3N2) infected Alveolus Chips (MOI = 1). (**f**) Volcano plot of DEGs in endothelial cells from HK/68 (H3N2)-infected Alveolus Chips (MOI = 1) compared to control uninfected chips. The names of DEGs from the S100 protein family are shown. (**g**) Gene Set Enrichment Analysis (GSEA) plots showing the significant enrichment of two gene sets in endothelial cells from HK/68 (H3N2)-infected (MOI = 1) compared with control uninfected Alveolus Chips. (**h**) Whole chip fluorescence imaging for CellTracker Green-labeled PBMCs at the endothelial cell surface 2 hours after perfusion through the vascular channel of uninfected control (Con) versus HK/68 (H3N2)-infected Alveolus Chips at 24 hpi (MOI = 1). (bar, 1 mm). (**i**) Higher magnification of images showing representative regions of Chips from (H) (bar, 100 µm). (**j**) Graph showing number of PBMCs recruited to the endothelium in response to infection by HK/68 (H3N2) (MOI = 1) and the baseline level of PBMCs in uninfected chips (Con). Data represent mean ± SD.; n = 3-4 biological replicates; unpaired two-tailed t-test, ***p<0.001.

As cytokine production by lung epithelial and endothelial cells is a key feature of early host innate immune response to viral infections, we analyzed cytokines that are secreted into the basal vascular outflow using Luminex assays, analogous to measurement of plasma cytokine levels in vivo. These studies revealed that infection with influenza H3N2 virus induces significantly higher levels of IL-6, IL-8, IP-10, TNF-α, and GM-CSF in the vascular outflows at 48 hours post infection (hpi) compared to control uninfected chips (**Fig. 2c**). RNA-seq of the epithelial cells carried out at the same time point (48 hpi) revealed 496 upregulated genes and 538 downregulated genes (**Fig. 2d**). GO analysis indicated that the differentially expressed genes (DEGs) belong to biological pathways including cell division and DNA replication that may be associated with antiviral defense responses and activation of alveolar barrier repair in response to virus induced injury (**Fig. 2e**). Some of these DEGs, such as CXCL10 and IL6, were further confirmed by qPCR (**Supplementary Fig. 5a**).

A more detailed analysis of antiviral responses induced in the lung epithelium in the Alveolus Chip revealed a dominant type III interferon (IFN-III) response mediated by interferon lambda (IFN λ) (**Fig. 2d and Supplementary Fig. 5b**). Importantly, this finding differs from results obtained from analysis of human lung alveolar epithelial cells cultured in static 2D culture on rigid planar dishes where IFNβ-mediated type I interferon response prevails^27^. However, our results replicate *in vivo* findings^28^, which suggest that IFN-III mediates the front-line response to viral infection in mucosal tissues^29^. In contrast to the epithelium (**Fig. 2d**), a more limited effect on the host transcriptome was observed in the endothelium (**Fig. 2f**). But enrichment analysis shows that genes involved in activation of immune responses and ion transport are also upregulated in these cells (**Fig. 2g**). Interestingly, however, this is likely through tissue-tissue signaling cross-talk as viral mRNA could not be detected within the endothelium.

Influenza virus infection also upregulates expression of endothelial adhesion molecules, thereby allowing the recruitment of leukocytes to the alveolus in vivo^11^. When fluorescently-labeled primary human peripheral blood mononuclear cells (PBMCs) were flowed through the endothelium-lined vascular channel 24 hpi with influenza H3N2, we detected upregulation of ICAM1 and TNFα in the endothelial cells by qPCR, indicating endothelial inflammation (**Supplementary Fig. 5c**). Indeed, this resulted in more than a 100-fold increase in the adhesion of the circulating PBMCs to the surface of the activated endothelium compared to control uninfected chips (**Fig. 2h-j and Supplementary Fig. 5d**). Thus, the human breathing Alveolus Chip faithfully replicates multiple system-level host innate immune responses of lung alveoli to influenza A virus infection.

### Breathing-like mechanical deformations suppress viral infection on-chip

We next explored the role of cyclic respiratory motions in virus-induced respiratory infections by infecting Alveolus Chips with H3N2 virus on day 15 of culture and measuring viral loads in the presence or absence of physiological, breathing-like, mechanical deformations (5% strain, 0.25 Hz). Immunostaining for influenza virus nucleoprotein (NP) revealed significant suppression of viral infection in lung alveolar epithelial cells exposed to breathing motions 2 days following introduction of virus on-chip compared to static chip controls (**Fig. 3a**). This inhibition by mechanical stimulation was further confirmed using qPCR and a plaque assay, which respectively show that that application of cyclic mechanical strain leads to a 50% reduction of viral mRNA in the epithelium (**Fig. 3b**) and ∼80% reduction of viral titers in apical washes compared to static controls (**Fig. 3c**). This also was accompanied by a reduction in production of the inflammatory cytokines, IP-10 and TRAIL, as measured in the vascular outflows from the chips (**Fig. 3d**). Moreover, similar inhibition of viral infection by cyclic mechanical strain was observed when the Alveolus Chips were infected with other respiratory viruses, including H5N1 influenza virus (**Fig. 3e**) and the common cold OC43 coronavirus (**Fig. 3f**).

**Figure 3.**
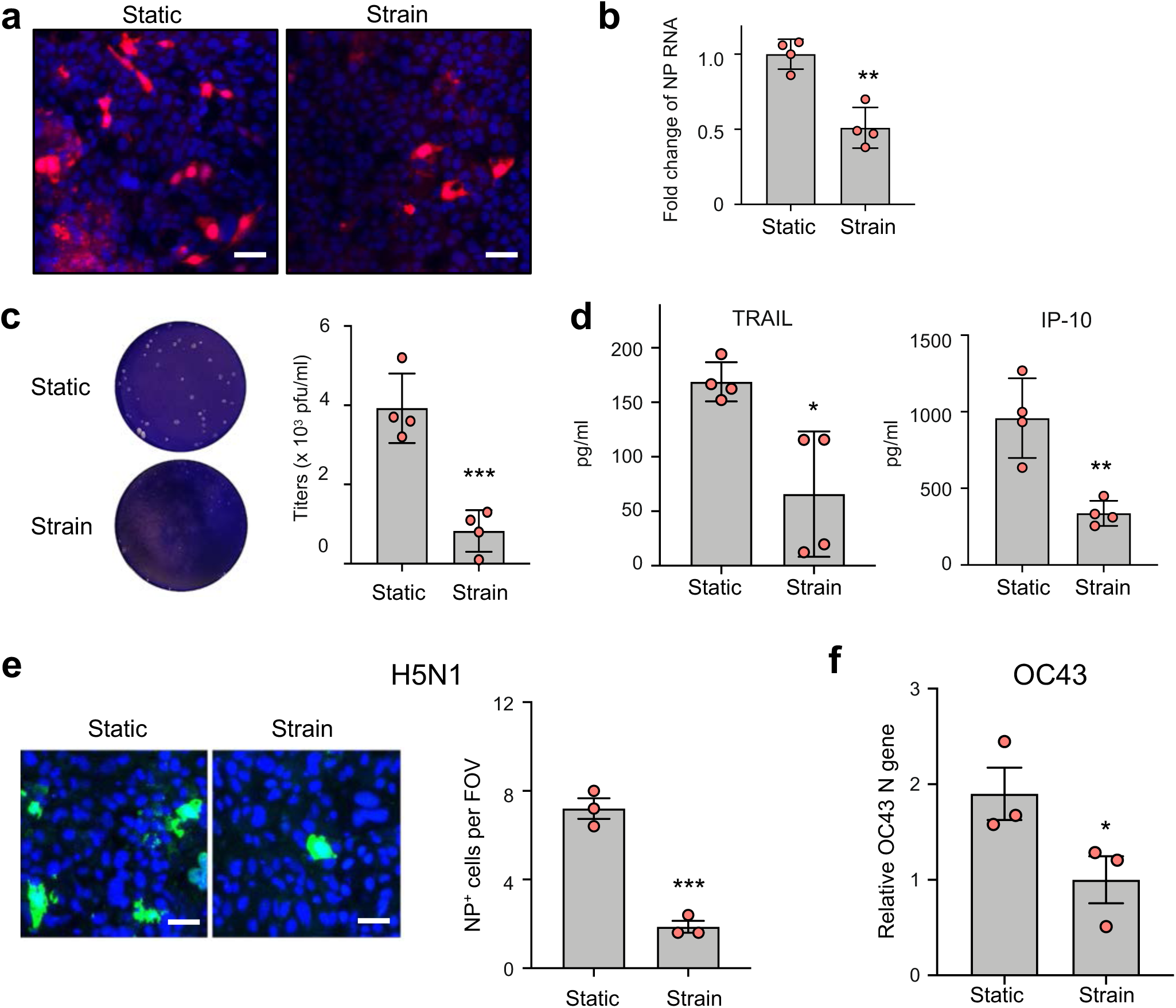
Cyclic mechanical strain inhibits viral infection in Alveolus Chip. (**a**) Immunostaining of influenza virus NP (red) in Alveolar Chips that are cultured under static condition (Static) or under 5% and 0.25 Hz cyclic mechanical strains (Strain) for 48 hours and then infected with HK/68 (H3N2) (MOI = 1) for another 48 hours (blue, DAPI-stained nuclei; bar, 50 µm). (**b**) Graph showing fold changes in RNA levels of influenza virus NP in the epithelium within Alveolus Chips from (A) as measured by qPCR. Data represent mean ± SD; n = 4 biological replicates; unpaired two-tailed t-test, **p<0.01. (**c**) Images (left) and graph (right) showing plaque titers of virus in the apical washes of Alveolus Chips from (A). Data represent mean ± SD; n = 4 biological replicates; unpaired two-tailed t-test, ***p<0.01. (**d**) Graphs showing cytokine production in the vascular effluents of Alveolus Chips from (A) at 48 hpi. Data represent mean ± SD; n = 4 biological replicates; unpaired two-tailed t-test, *p<0.05, **p<0.01, ***p<0.001. (**e**) Immunofluorescence micrographs (left) showing influenza virus NP (green) in Alveolar Chips that are cultured under static condition (Static) or under 5% and 0.25 Hz cyclic mechanical strains (Strain) for 48 hours and then infected with H5N1 (MOI = 0.001) for another 48 hours (blue, DAPI-stained nuclei; bar, 50 µm), and a graph (right) showing the numbers of NP^+^ cells per field of view (FOV). Data represent mean ± SD; n = 3 biological replicates; unpaired two-tailed t-test, ***p<0.001. (**f**) Graph showing relative levels of OC43 N gene measured by qPCR in epithelium of Alveolar Chips that are cultured under static condition (Static) or under 5% and 0.25 Hz cyclic mechanical strains (Strain) for 48 hours and then infected with OC43 (MOI = 5) for another 48 hours. Data represent mean ± SD; n = 3 biological replicates; unpaired two-tailed t-test, *p<0.05.

### Mechanical strain increases innate immunity in Lung Chips

To gain insight into how mechanical strain associated with physiological breathing motions might normally act to combat viral infection, we analyzed the RNA-seq analysis time course and noticed that exposure to continuous cyclic mechanical deformations resulted in increased expression of multiple IFN-related antiviral genes, including DDX58, MX1, OAS1, and STAT1, in both lung epithelial and endothelial cells from day 8 to day 14 of culture (**Fig. 1h and Supplementary Fig. 4b**). Indeed, many interferon-stimulated genes (ISGs) have higher expression in mechanically stimulated cells at day 14 on-chip but not in cells cultured in parallel in static Transwells (**Supplementary Fig. 6a**).

To directly determine whether mechanical strain influences innate immunity pathway signaling, we cultured the Alveolus Chips in the presence or absence of cyclic mechanical strain from day 10 to 14 and performed RNA-seq analysis. Consistent with our earlier results, lung alveolar epithelial cells mechanically stimulated on-chip exhibit higher expression of ISGs and cytokines, such as MX1, MX2, IL-6, CXCL10, CXCL5, IFI44L, and IFIH1, compared to static chip controls (**Fig. 4a**). Functional enrichment analysis of DEGs demonstrated that application of physiological cyclic strain activates pathways related to host defense response, while suppressing processes related to cell cycle and cell proliferation (**Fig. 4b**).

**Figure 4.**
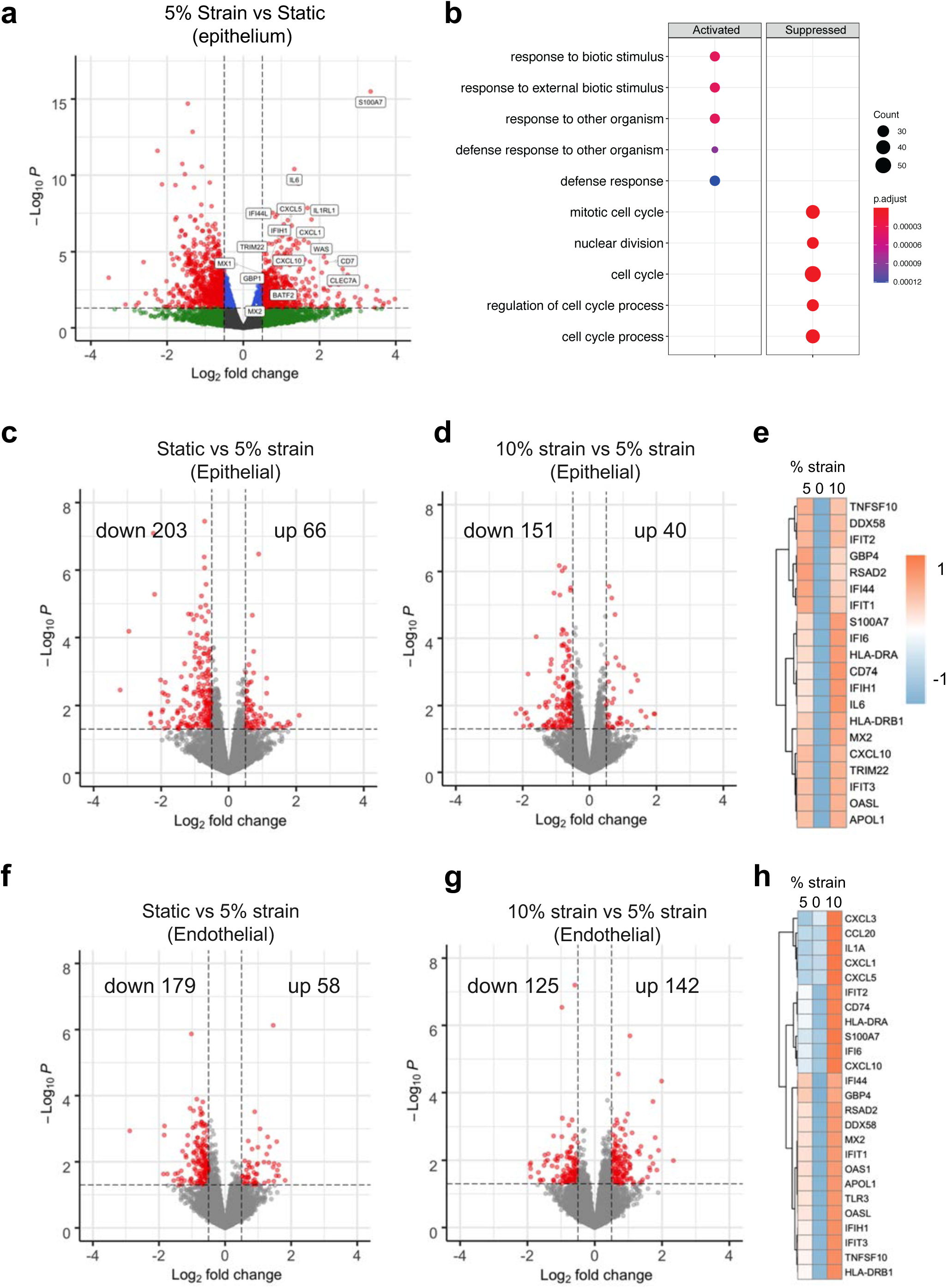
Mechanical strain reversibly regulates innate immune response in Alveolus Chip. (**a**) Volcano plot of DEGs comparing epithelial cells from Alveolus Chips under static or 5% strain culture condition for 4 days. DEGs (P_adj_ < 0.05) with a fold change >1.5 (or <−1.5) are indicated in red. The names of DEGs belonging to the innate immune pathway are labeled. (**b**) Dot plot showing the biological processes activated or suppressed by 5% strain vs. static culture condition in Alveolus Chips from (A). (**c & d**) Volcano plots of DEGs showing the effects of switching 5% strain to static for 2 days on epithelial cells on-chip (**c**) and the effects of increasing 5% strain to 10% strain for 2 days on epithelial cells on-chip (**d**). (**e**) Heat map showing differentially expressed innate immune genes in epithelial cells under different magnitudes of mechanical strains. (**f & g**) Volcano plots of DEGs showing the effects of switching 5% strain to static for 2 days on endothelial cells on-chip (**f**) and the effects of increasing 5% strain to 10% strain for 2 days on endothelial cells on-chip (**g**). (**h**) Heat map showing differentially expressed innate immune genes in endothelial cells under different magnitudes of mechanical strains.

To further investigate the association between mechanical strain and innate immunity, we switched a subset of chips that had been exposed to 5% cyclic mechanical strain from day 10 to 14 to static conditions or to elevated mechanical strain (10%) condition on day 15 while continuing to mechanically stimulate the remaining chips at 5% for 2 additional days. RNA-seq analysis of the epithelial cells on day 17 revealed that cessation of mechanical stimulation results in suppression of the innate immune response, exemplified by decreased expression of antiviral genes, such as IFIT1, IFIT2, OASL, and DDX58 (**Fig. 4c and Supplementary Fig. 6b)**. On the other hand, elevating the level of mechanical stimulation to 10% strain results in modest upregulation of innate immune response genes (**Fig. 4d,e** and **Supplementary Fig. 6c**). In parallel, similar reduction in innate immune response pathway signaling is observed in endothelial cells when mechanical stimulation ceases (**Fig. 4f and Supplementary Fig. 6d**). However, increasing mechanical strain to 10% results in further upregulation of innate immune response genes (**Fig. 4g, h** and **Supplementary Fig. 6e**), including several inflammatory cytokines, such as CXCL3, CCL20, IL1A, CXCL1, and CXCL5 (**Fig. 4h**). These results suggest that endothelial cells may be more sensitive to elevated mechanical strains than the epithelial cells. Together, these results demonstrate that cyclic mechanical forces similar to those experienced during breathing in the lung sustain higher levels of antiviral innate immunity in both lung alveolar epithelial cells and pulmonary microvascular endothelial cells compared cells cultured in the same chips under static conditions, which helps to explain why viral infection efficiency is reduced in the mechanically strained Alveolus Chips.

### S100A7 and RAGE mediate the effects of mechanical strain on lung innate immunity

We then leveraged this Organ Chip approach to explore the underlying mechanism responsible for these effects by examining our RNA-seq data sets. This analysis revealed that the S100 calcium-binding protein A7 (S100A7), a member of the S100 family of alarmins proteins, as one of the top genes differentially expressed in response to cyclic strain application in all experiments (**Fig. 1h, Fig. 4 a,c,e**, and **Supplementary Fig. 7a**), and one that was not induced cells in static culture on-chip or in Transwells (**Fig. 5a and Supplementary Fig. 7a**).

**Figure 5.**
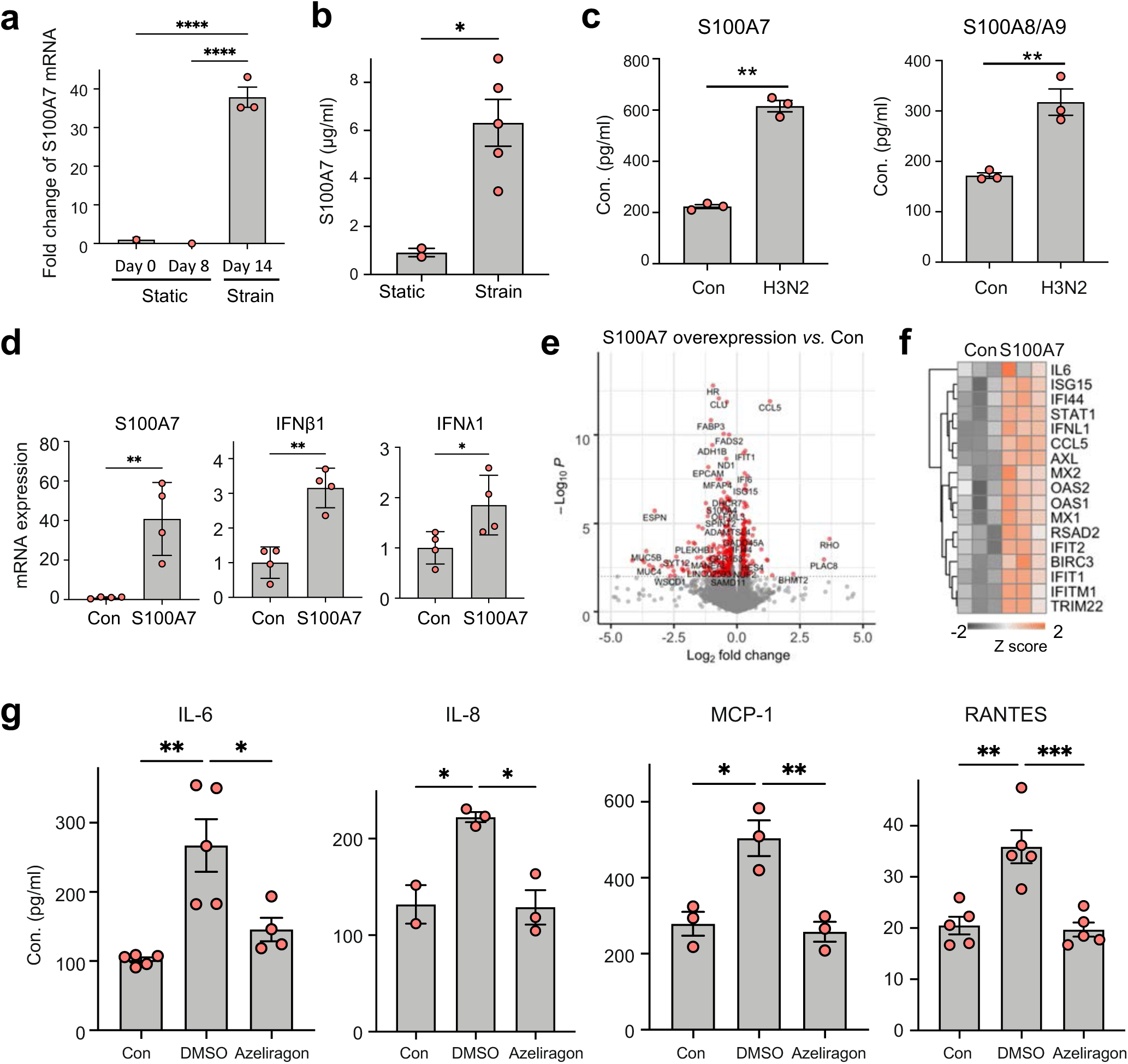
Mechanical strain-induced S100A7 increases innate immunity. (**a**) Graph showing the levels of S100A7 mRNA (transcripts per million) in the epithelial cells of Alveolus Chips at different time points of culture, measured by RNA-seq. Data represent mean ± SD; n = 3 biological replicates; one-way ANOVA with Tukey’s multiple comparisons test, ****p<0.0001. (**b**) Graph showing the protein levels of S100A7 in the apical washes of Alveolus Chips under strain or static condition, measure by ELISA. Data are shown as mean ± SEM; n =2-5 biological replicates; unpaired two-tailed t-test, *p<0.05. (**c**) Graph showing the protein levels of S100A7 (left) and S100A8/A9 (right) in the vascular effluents of the Alveolus Chips at 48 hpi with HK/68 (H3N2) virus (MOI = 1). Data represent mean ± SD; n = 3 biological replicates; unpaired two- tailed t-test, **p<0.01. (**d**) Graphs showing fold changes in mRNA levels of S100A7, IFNβ1, and IFNλ1 in epithelial cells of the Alveolus Chips that were transfected with human S100A7-expressing plasmid or the vector control (Con) for 48 hours. Data are shown as mean ± SD; n = 4 biological replicates; unpaired two-tailed t-test, *p<0.05, **p<0.01. (**e**) Volcano plots of DEGs showing the effects of S100A7 overexpression on transcriptome in epithelial cells of the Alveolus Chips that were transfected with S100A7-expressing plasmid for 2 days with the empty plasmid as a control (Con). (**f**) Heat map showing that S100A7 upregulates the expression of many genes involved in innate immune response in epithelial cells of the Alveolus Chips. n = 3 biological replicates. (**g**) Graphs showing the levels of cytokines in the vascular effluents of Alveolus Chips that were uninfected (Con), or infected with HK/68 (H3N2) (MOI = 1) in the presence or absence of 100 nM Azeliragon. Data are shown as mean ± SD; n = 2-5 biological replicates; one-way ANOVA with Tukey’s multiple comparisons test, *p<0.05, **p<0.01, ***p<0.001.

This increase in S100A7 gene expression in response to mechanical stimulation is also associated with a significant increase in S100A7 protein secretion, as determined by quantifying its levels in apical washes from human Alveolus Chips using an enzyme-linked immunosorbent assay (ELISA) (**Fig. 5b**). Interestingly, S100A7 gene expression and protein secretion are also upregulated in both epithelial cells and endothelial cells in response to infection of the Alveolus Chips with H3N2 influenza virus, and this is accompanied by upregulation of other S100 family members, including S100A8, S100A9, and S100A12 (**Fig. 2f** and **Fig. 5c**). Moreover, over- expression of S100A7 is sufficient to induce significant upregulation of the antiviral cytokines IFNβ1 and IFNλ1 in lung alveolar epithelial cells on-chip (**Fig. 5d**) as well as in conventional monoculture (**Supplementary Fig. 7b**) and in A549 lung cells, which similarly show both overexpression of S100A7 and reduced influenza H1N1 virus infection (**Supplementary Fig. 7c,d**). To directly examine the effect of S100A7 on gene transcriptional network, we performed RNA-seq analysis of alveolar epithelial cells on-chip after transfection of a plasmid expressing human S100A7. Differential gene expression analysis revealed that increased expression of S100A7 indeed upregulates many genes involved in the innate immune response, including IFNL1, CCL5, IFI6, and IL-6 (**Fig. 5e, f**). Therefore, S100A7 that is induced by breathing motions is sufficient to drive the host innate immune response.

A role for S100A7 in antiviral host responses has not been reported previously; however, it attracted our interest because it has been shown to exert immunomodulatory effects by binding to the inflammation associated Receptor for Advanced Glycation End Products (RAGE)^30, 31^. Importantly, RAGE deficiency increases sensitivity to viral infection in animal models^32^, and the closely related S100A8 and S100A9 proteins have been shown to induce innate immune programming and protect newborn infants from sepsis^33^. S100A7 also has been shown to exhibit antibacterial and antifungal activities^34, 35^. However, while these S100 proteins help to resolve infections, they can cause severe inflammation and tissue damage when they are aberrantly expressed^36^. For example, S100A8, S100A9, and S100A12 levels in blood have recently been shown to correlate with disease severity in COVID-19 patients infected with SARS-CoV-2^37, 38^.

As S100A7 and other members of the S100 family signal through RAGE^39^, we next explored whether RAGE inhibitors can be used to suppress host inflammatory responses during viral infection in Lung Alveolus Chips. One advantage of human Organ Chips is that they permit drug testing in a more clinically relevant pharmacological setting by enabling clinically relevant doses of drugs (e.g., maximum blood concentration or C_max_) to be perfused through the vascular channel under flow as occurs in the vasculature of living organs^40^. When azeliragon (TTP488), a RAGE inhibitor that is currently in Phase III clinical trials for patients with Alzheimer Disease, is perfused at its C_max_ through the vascular channels of mechanically strained human Alveolus Chips infected with H3N2 influenza virus, it significantly blocks induction of multiple cytokines, including IL-6, IL-8, IP-10 and RANTES (**Fig. 5g**), and similar effects are produced using a different RAGE inhibitor drug (FPS-ZM1) (**Supplementary Fig. 7e**). Thus, the effects of mechanical breathing motions on host innate immune immunity in human Lung Alveolus Chips appear to be mediated by modulation of signaling through the S100A7-RAGE pathway.

## DISCUSSION

Here we describe how use of human Organ Chip technology has enabled discovery of a novel mechano-immunological control mechanism in which physiological breathing motions suppress viral infection in human lung alveoli by activating host innate immune responses.

Exposure to cyclic mechanical cues associated with physiological breathing motions (5% strain; 15 deformations/min) sustains moderate levels of production and secretion of S100A7 protein by alveolar epithelium and endothelium, which binds to RAGE and generates an innate immune response that protects against viral infection, whereas S100A7 levels are lower and viral infection greater when breathing motions are absent. Our data also show that pharmacological inhibitors of RAGE signaling suppress inflammation during influenza infection, raising the possibility that RAGE inhibitors might represent novel adjuvants that may be used in combination with antiviral therapies to suppress life-threatening inflammation of the distal airway that can be triggered by influenza A viruses. Importantly, they might also be useful for suppressing augmented inflammatory responses associated with conditions, such as COVID-19, which result in abnormally high levels of S100 family protein production^37, 38^.

Lung alveoli are exposed to dynamic mechanical stresses with each breath and these physical deformations are known to influence lung development^2, 41, 42^, surfactant production^42, 43^, and tissue barrier integrity^4^. Mechanical forces also have been suggested to activate immune cells in lung^44^, particularly in response to pathological stresses associated with hypeventilation^7, 45^. But little is known about the effects of breathing-associated cyclic mechanical strain on host innate immunity in response to viral infections within the tissue parenchyma. Our study provides the first evidence indicating that physical forces control innate immunity in nonimmune cells and tissues, specifically in pulmonary alveolar epithelium and microvascular endothelium. This is important for the lung as it is exposed to numerous airborne antigens during early life while the adaptive immune system is still immature^46^. These findings are also supported by recent single cell RNA-seq studies of lung alveolar epithelium, which demonstrate increased innate immunity at birth in both mouse and human^20, 21^. In addition, our finding that a hyper-physiological deforming stress (10% strain) leads to an augmented host inflammatory response, especially in the endothelial cells, resonates with clinical findings, which show that low tidal volume ventilation that induces less cyclic mechanical strain results in lower plasma levels of inflammatory cytokines than use of a normal tidal volume strategy^47, 48^; importantly, this greatly benefits patients with acute respiratory distress syndrome^49^. Thus, human Alveolar Chips can be used in the future to apply strain deformations at higher amplitudes and at higher frequencies to better determine the parameter space in relation to lung innate immunity, and thereby help to provide design criteria to optimize therapeutic strategies to mitigate ventilator-associated lung injury.

This work led to the discovery that production of S100A7 protein and its binding to RAGE mediate the mechanical induction of innate immunity we observed in the Alveolar Chips. S100A7 has been previously shown to exerts its immunomodulatory functions via RAGE-dependent activation of MAPK and NFκB signaling pathways^31^. Interestingly, RAGE is more highly expressed in lung ATI cells at baseline than any other cell type, many of which show little to no RAGE expression^50^, and it appears to play a major role in pulmonary inflammation in various diseases, including COVID-19^50, 51^. Our finding that S100A7 expression is controlled mechanically in lung alveoli is also consistent with the observation that it is also induced by mechanical stress in human dental pulp cells^52^. Activation of the S100A7-RAGE pathway may be essential for development of host innate immunity in newborns as well since high amounts of the perinatal alarmins S100A8 and S100A9 have been reported to induce innate immune programming and protect newborn infants from sepsis^33^. Understanding the precise molecular signaling mechanism by which mechanical forces induce S100A7 is beyond the scope of the present study, but it could involve activation of mechanosensitive ion channels such as Piezo1 or TRPV4, which have been shown to mediate activation of immune cells in lung^7, 44, 45^.

Finally, we demonstrated that influenza virus infection also induces the expression of S100A7, S100A8, S100A9, and S100A12 in the Alveolus Chip, a finding that is consistent with recent clinical observations from COVID-19 patients that exhibit late stage lung infections with SARS-CoV-2 virus^37, 38, 53–56^. While the innate immune response mediated by the S100 alarmins may be beneficial in combating against initial infection, sustained expression and dysregulation of alarmins can result in hyperinflammatory responses that cause irreversible tissue damage, as encountered by patients with severe diseases that require mechanical ventilation^57^. Importantly, using a clinical-relevant dosing strategy in the Alveolus Chips (i.e., by perfusing them with drugs at their clinical C_max_), we found that administration of RAGE inhibitor drugs, such as azeliragon and FPS-ZM1, inhibit viral-induced secretion of inflammatory cytokines. The translational potential of RAGE inhibitors awaits further testing in animal models. Nevertheless, it represents an attractive treatment option for attenuating aberrant host immune responses due to viral infection and/or mechanical ventilation in patients with pneumonia or acute respiratory distress syndrome.

As demonstrated by the current COVID-19 pandemic, potential pandemic viruses, including influenza A viruses, pose a major threat to public health. The lack of experimental models of the distal lung has greatly handicapped our efforts to repurpose existing drugs, understand disease pathology, and develop new therapies. Using influenza A virus as an example, we extend past work focused on viral infection of large airway^26^ by demonstrating that the Alveolus Chip recapitulates viral receptor expression, strain-dependent infectivity, and system-level host responses in the alveolar epithelial, endothelial, and immune cells during infection by viruses that target alveoli. These features have been overlooked in previous influenza research using cell lines or animal models, but are indispensable for better understanding of disease pathogenesis and identification of more predictive biomarkers and effective therapeutic targets. Thus, in addition to serving as a model system for gaining greater insight into mechanochemical mechanisms that underlie this form of innate immunity, it might serve as a useful preclinical model for identification and optimization of new and more effective antiviral therapeutics for diseases of the distal airway that lead to alveolitis and acute respiratory distress syndrome.

## Acknowledgments

This research was supported by the Wyss Institute for Biologically Inspire Engineering at Harvard University, Defense Advanced Research Projects Agency (DARPA) under Cooperative Agreement (HR00111920008 and HR0011-20-2-0040), and NIH (UH3-HL-141797). We thank Genewiz Inc. for providing RNA-seq. We thank Maurice Perez, Michael Carr, Eric Zigon and Thomas Ferrante for their assistance in the lab support and facility usage.

## Author Contributions

H.B., L.S., R.PB. and D.E.I. conceived this study. H.B. and L.S. designed experiments and analyzed data. H.B., L.S., A.J., C.B., R.P., C.O., and M.R., and A.N., and performed experiments. R.K.P. assisted with bioinformatics analysis. S.G., G. G., and R.P. assisted in experiments design and data analysis. H.B., L.S., and D.E.I. wrote the manuscript with input from other authors.

## METHODS

### Alveolus Chip Culture

Microfluidic two-channel Organ Chip devices and automated ZOE→ instruments used to culture them were purchased from Emulate Inc (Boston, MA, USA). After chip activation using ER1/ER2 reagents following manufacturer’s instruction, both channels were seeded with 200 µg/ml Collagen IV (5022-5MG, Advanced Biomatrix) and 15 µg/ml of laminin (L4544-100UL, Sigma) at 37°C overnight. The next day (day 1), primary human lung microvascular endothelial cells (Lonza, CC-2527, P5) and primary human lung alveolar epithelial cells (Cell Biologics, H- 6053) were sequentially seeded in the bottom and top channels of the chip at a density of 8 x 10^6^ and 1.6 x 10^6^ cells/ml, respectively, under static conditions. On day 2, the chips were inserted into Pods→ (Emulate Inc.), placed within the ZOE→ instrument, and the apical and basal channels were respectively perfused (60 µL/hr) with epithelial growth medium (Cell Biologics, H6621) and endothelial growth medium (Lonza, CC-3202). On day 5, 1 µM dexamethasone was added to the apical medium to enhance barrier function. On day 7, an air-liquid interface (ALI) was established in the epithelial channel by flowing at 1,000 μL/hr for 5 minutes until all medium from this channel was emptied. Chips were fed through the lower vascular channel, and this medium was changed to EGM-2MV with 0.5% FBS on day 9. Two days later, the ZOE→ instrument was used to apply cyclic (0.25 Hz) 5% mechanical strain to the engineered alveolar-capillary interface to mimic lung breathing on-chip. Chips were used for experiments on Day 15.

### Viral stocks

Influenza virus strains used in this study include A/WSN/33 (H1N1), A/Hong Kong/8/68/ (H3N2), and A/Hong Kong/156/1997 (H5N1). All viruses were obtained from the Centers for Disease Control and Prevention (CDC) or kindly shared by Drs. P. Palese, R.A.M. Fouchier, and A. Carcia-Sastre. Influenza virus strains were expanded in MDCK.2 cell^58^. OC43 coronavirus (VR-1558) was obtained from the ATCC and expanded in HCT-8 cells (ATCC) as previously described ^59^.

### Viral infection on chip

Chips and pods were removed from ZOE→ and put in biosafety cabinet (BSC). Chips were disconnected from pods and 40 µl viral inoculate was added to the top channel inlet to infect alveolar epithelial cells. Then chips were put back to pods and kept in static condition at 37 °C. 2 hours later, chips were removed from pods and top channels were washed with 100 µl DPBS (-/-). Chips were reprimed, connected to pods and continued for flow at 60 µL/hr and 5% mechanical strain.

### RNA extraction from Chip for RNA-seq

For RNA extraction, chips were removed from pods. An empty 200 µL filtered tip was inserted to top channel outlet and washed with 100 µL DPBS (+/+) into the top channel inlet to wash the apical channel. After collecting top channel washes, a new empty 200 µL filtered tip was inserted to top channel outlet and 100 ul RNase easy lysis buffer (Qiagen, #74034) was used to lyse epithelial cells by quickly pressing and releasing the micropipette plunger 3 times. Lysates were collected in a clean labelled 1.5 ml tube and stored at -80°C for RNA-seq analysis. Endothelial cell lysates were subsequently collected in a similar manner.

### RNA-seq and bioinformatic analysis

RNA-seq was performed by Genewiz using a standard RNA-seq package that includes polyA selection and sequencing on an Illumina HiSeq with 150-bp pair-ended reads. Sequence reads were trimmed to remove possible adapter sequences and nucleotides with poor quality using Trimmomatic v.0.36. The trimmed reads were mapped to the Homo sapiens GRCh38 reference genome using the STAR aligner v.2.5.2b. Unique gene hit counts were calculated by using feature Counts from the Subread package v.1.5.2 followed by differential expression analysis using DESeq2. Gene Ontology analysis was performed using DAVID^60^ and the Enrichplot R package. Volcano plots were generated using the EnhancedVolcano R package. Heatmaps and scatter plots were generated using the ggplot2 R package. STEM ^16^ was applied to identify significant temporal patterns using the expression profiles of differentially expressed genes. Five significant patterns were identified with p-value < 0.01 and cluster size >300 genes. RNA-sequencing data have been deposited in the Gene Expression Omnibus (GEO) database under the accession code xxx.

### RT-qPCR

Total RNA was isolated using the RNeasy Mini plus Kit or the RNeasy Micro plus Kit (Qiagen). After determining RNA concentrations by spectrophotometry, 50 to 500 ng of total RNA was used for cDNA synthesis. Reverse transcription was conducted using the Omniscript RT Kit (Qiagen) or the Sensiscript RT kit (Qiagen). Quantitative real-time PCR was performed using the SsoAdvanced- Universal SYBR Green Supermix (Biorad). The specificity of primers was confirmed by melting curve analysis and gel electrophoresis. qPCR was performed on a CFX Connect Real Time PCR Detection System (Biorad). Relative RNA level was quantified using the ΔΔCt method^61^ and normalized to the endogenous control GAPDH unless specified otherwise. All primers were purchased from IDT (supplementary table 1).

### PBMC study on chip

At 23 hours after infection, PBMC was prepared for the adhesion study. PBMC (StemCell, #70025.1) was thawed and resuspend in 10 ml DMEM with 10% FBS and 1% Penicillin Streptomycin. 10 µl cell tracker green at a final concentration of 10 µM (Invitrogen, #C7025) was used to label cells for 20 min at 37 °C. Then cells were centrifuged and resuspend to 5 x 10^7^/ml. At 24 hours after infection, chips were detached from pods and 25 µl labeled PBMC were added to the bottom channel. Chips were flipped and placed on chip cradles. PBMC were allowed to adhere at 37 °C for 2 hours before flipped back and washed with 100 µl flow media to wash out unattached cells. Chips were immediately imaged under a fluorescent microscope.

### Immunostaining and confocal microscopy

Cells were rinsed with PBS(-/-), fixed with 4% paraformaldehyde (Alfa Aesar) for 20 min at room temperature, permeabilized with 0.1% Triton X-100 (Sigma-Aldrich) in PBS (PBSX) for 10 min, blocked with 5% goat serum (Life Technologies, #50062Z) in PBSX for 1 h at room temperature, and incubated with antibody diluted in blocking buffer (5% goat serum in PBSX) overnight at 4 °C, followed by incubation with fluorescent-conjugated secondary antibody for 1 h at room temperature; nuclei were stained with DAPI (Invitrogen) after secondary antibody staining. Fluorescence imaging was conducted using a confocal laser-scanning microscope (SP5 X MP DMI-6000, Germany) and image processing was done using the ImageJ software. Antibody

### Cytokine analysis

Vascular effluents from Alveolus Chips were collected and analyzed for a panel of cytokines and chemokines, including IL-6, IL-8, IP-10, TNF-α, RANTES, S100A8/A9, and GM-CSF using custom ProcartaPlex assay kits (Invitrogen). Analyte concentrations were determined using a Luminex100/200 Flexmap3D instrument coupled with Luminex XPONENT software (Luminex, USA). S100A7 was measured using an ELISA kit (Aviva Systems Biology) according to manufacturer’s instruction.

### RAGE inhibitor treatment on-chip

Azeliragon (TTP-488) and FPS-ZM1 were purchased from MedChemExpress (#HY-50682) and Sigma (#553030), respectively. Drugs were diluted using Alveolar Chip flow media to its Cmax (100 nM for Azeliragon and 200 nM for FPS-ZM1) with a final DMSO concentration of 0.5% and perfused at 60 µl/h through the vascular channel at 24 hours before infection with influenza virus.

### Culture and transfection of A549 cells

A549 cells (ATCC CCL-185) were cultured in Dulbecco’s modified Eagle’s medium (DMEM) (Life Technologies) supplemented with 10% fetal bovine serum (FBS) (Life Technologies) and penicillin-streptomycin (Life Technologies). All cells were maintained at 37 °C and 5% CO_2_ in a humidified incubator. Transfection was performed using TransIT-X2 Dynamic Delivery System (Mirus) according to the manufacturer’s instructions. pCMV6-XL5 empty plasmid (PCMV6XL5) and S100A7 plasmid were purchased from Origene (#SC122639). RAGE plasmid was purchased from Sino Biological (HG11629-ACG). Endotoxin-free plasmid purification was performed by Genewiz.

### Culture and transfection of human primary alveolar epithelial cells

Primary human lung alveolar epithelial cells (Cell Biologics, H-6053) was seeded on 0.1% gelatin-coated 12-well plate according to manufacturer’s instruction. When reaching 70% confluent, cells were transfected with pCMV6-XL5 empty plasmid or S100A7 plasmid using TransIT-X2 Dynamic Delivery System (Mirus).

### Transfection in Human Alveolus Chip

Human Alveolus Chips were transfected with plasmid DNA using the TransIT-X2 reagent (Mirus Bio). 3 ug of plasmid DNA and 6 ul TransIT-X2 reagent were constituted in 300 ul OPT-MEM before added to 3 ml chip flow medium. Then 1.5 ml of the solution was added to apical and basal inlet reservoirs and flew at 60 ul/h for 24 hours. Then apical channel was empties to reestablish ALI; basal channel was changed to normal flow medium for additional 24 hours before qPCR analysis.

### Plaque assay

Virus titers were determined by plaque assay. Confluent MDCK cell monolayers in 12-well plate were washed with PBS, inoculated with 1 mL of 10-fold serial dilutions of influenza virus samples for 1 hour at 37℃, and then overlaid with 1 mL of DMEM (Gibco) supplemented with 1.5% low melting point agarose (Sigma-Aldrich) and 2 µg/mL TPCK-treated trypsin (Sigma-Aldrich). 2-4 days after incubation at 37℃ under 5% CO_2_, the cells were fixed with 4% paraformaldehyde (Alfa Aesar) and stained with crystal violet (Sigma-Aldrich) to visualize the plaques; virus titers were determined as plaque-forming units per milliliter (PFU/mL).

### Statistics

Each experiment was repeated at least three times, with at least 3 biological replicates for each data point. Data are displayed as mean values ± standard deviation (SD) unless otherwise noted in the figure legends. Graphing and statistical comparison of the data were performed using Prism 9 (GraphPad Software). Unless otherwise noted, two-group comparisons were assessed using the two-tailed Student’s t test; comparison of three or more groups were analyzed by one-way ANOVA with Bonferroni multiple comparisons. P values less than 0.05 were considered to be statistically significant (*, P < 0.05; **, P < 0.01; ***, P < 0.001; n.s., not significant).

---

**Figure S1.**
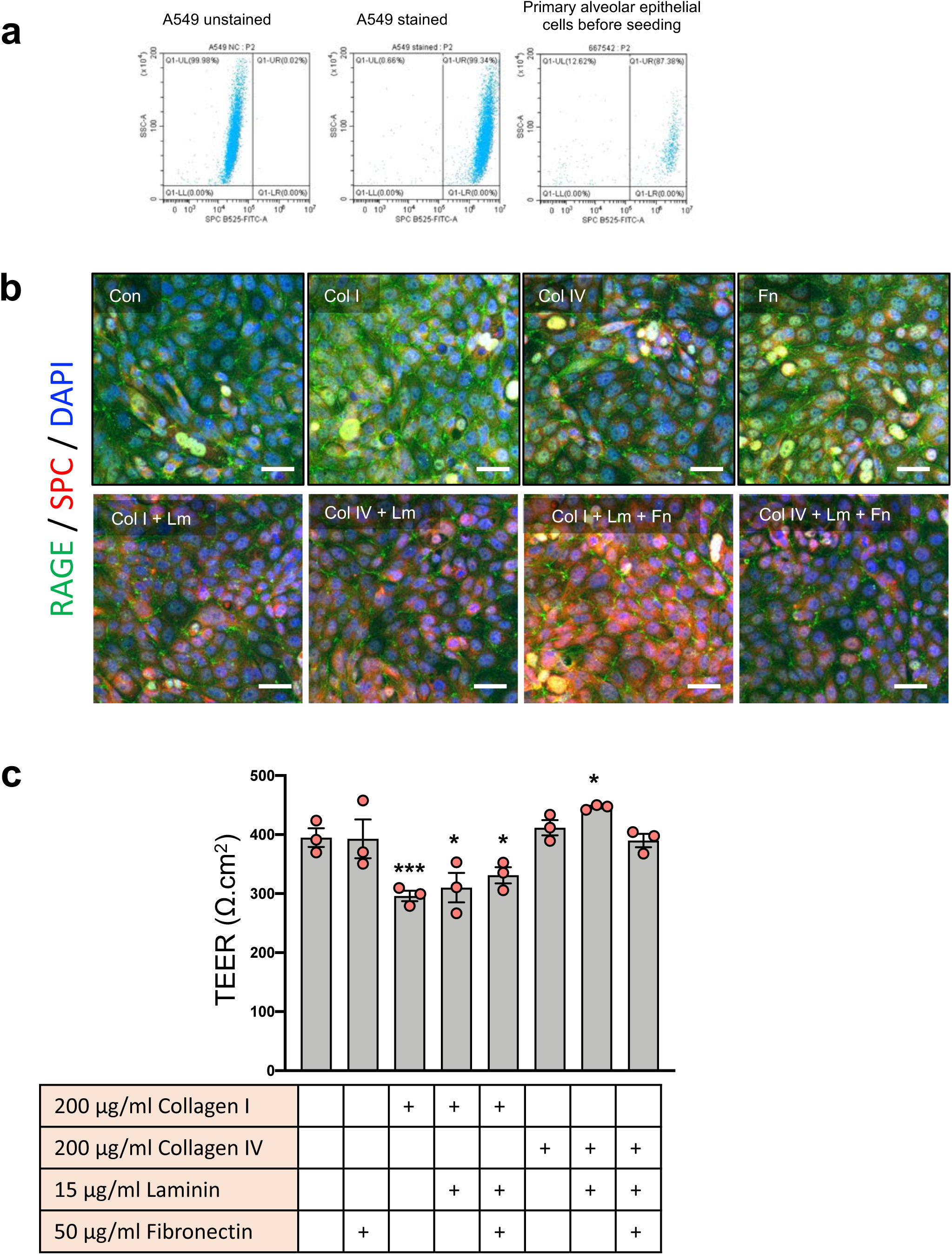
Protocol optimization for culture of human Alveolus Chip. (**a**) Flow cytometry showing 87% cells seeded onto chip are positive stained with type II marker SPC. A549 was used as a positive control for staining. (**b**) Immunofluorescence micrographs showing the expression levels and distribution of ATI cell marker RAGE and ATII cell marker SPC type II in the epithelium of Alveolus Chips coated with different ECM components. Con, without ECM; Col I, Collagen I; Col IV, Collagen IV; Fn, Fibronectin; Lm, Laminin. (bar, 50 μm) (**c**) The effects of different ECM coatings on barrier function as measured by transepithelial electrical resistance (TEER). Data are shown as mean ± SD; n = 3 biological replicates; unpaired two-tailed t-test (compared with no coating control), *p<0.05, ***p<0.001.

**Figure S2.**
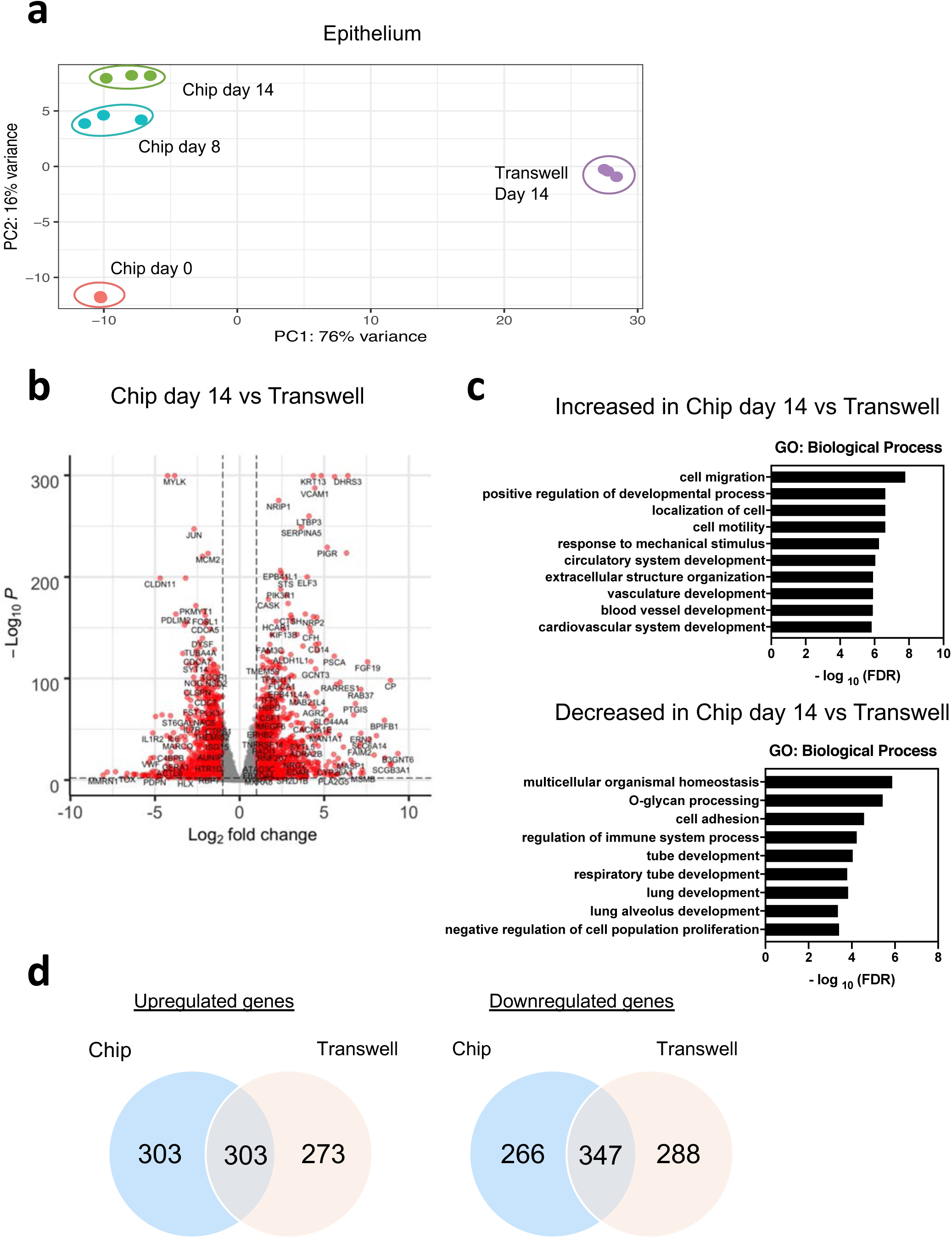
Characterization of Alveolus Chips and comparison with Transwell culture. (**a**) PCA plot of RNA-seq datasets from the Alveolus Chip or the transwell. (**b**) Volcano plot of DEGs comparing epithelial cells from Alveolus Chips at day 14 of culture with Transwell culture. DEGs (P_adj_ < 0.01) with a fold change >2 (or <−2) are indicated in red. The names of top DEGs are shown. (**c**) Gene ontology analysis showing enriched biological processes from (**b**). (**d**) Venn diagram to show the numbers of common and unique gene regulation in epithelial cells of Alveolus Chip and the Transwell culture.

**Figure S3.**
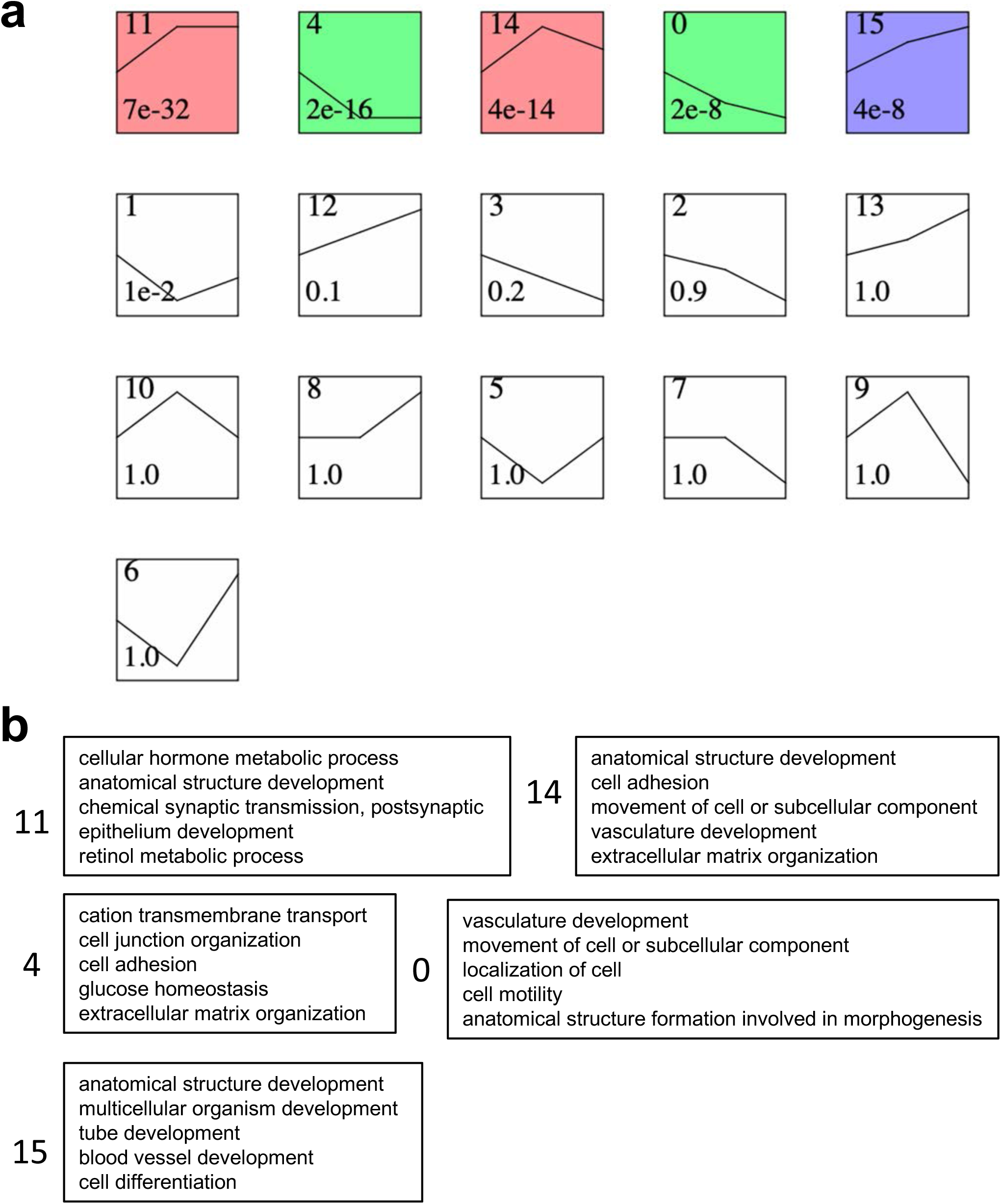
Temporal gene expression patterns in epithelial cells of Alveolus Chip. (**a**) All gene expression patterns identified using short time-series expression miner (STEM). Significant patterns were colored and p value was shown inside the box. (**b**) Functional annotations enriched by the temporal gene expression patterns that are significant in (**a**).

**Figure S4.**
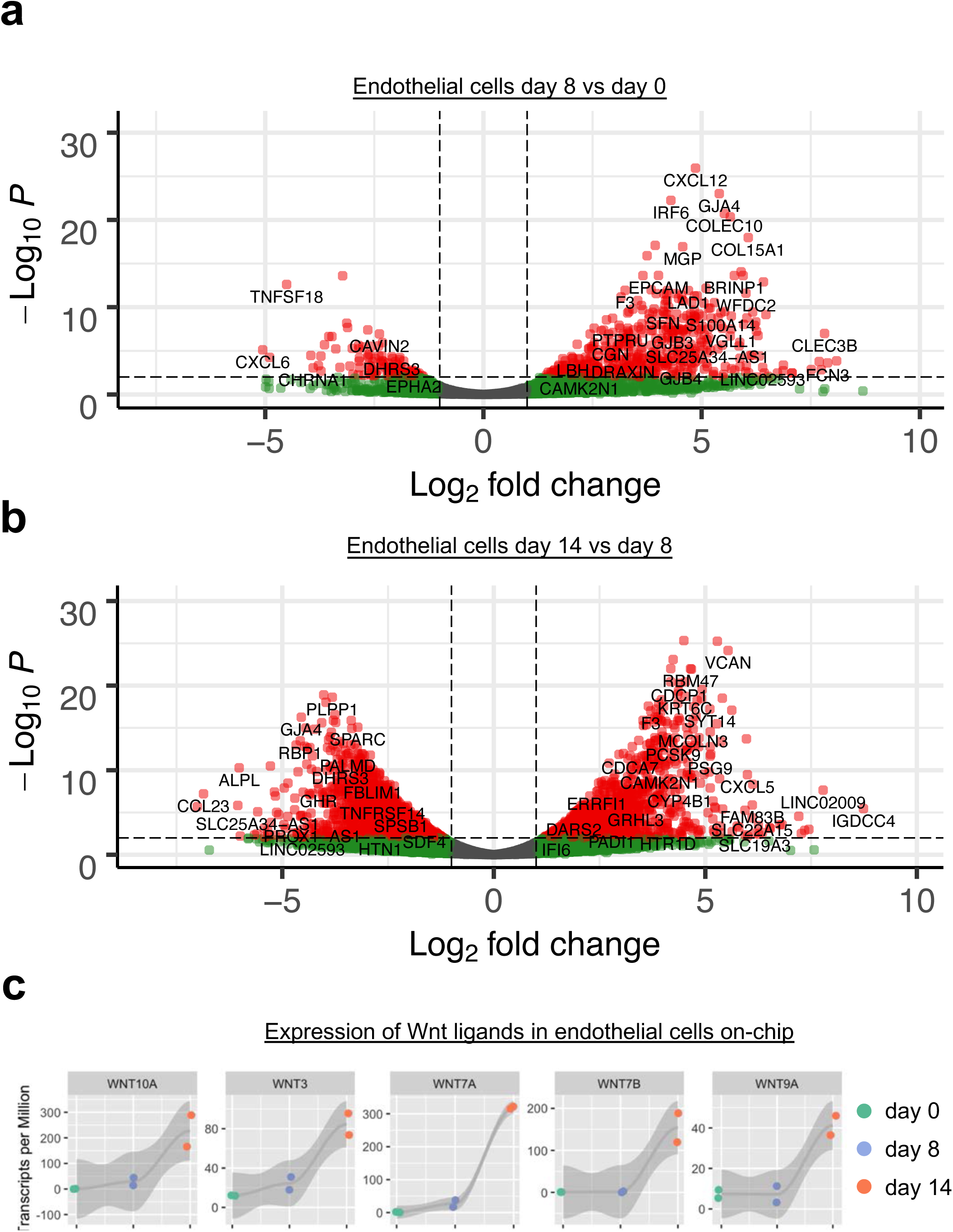
Transcriptomic changes in endothelial cells of Alveolus Chip. (**a**) Volcano plot of DEGs showing the transcriptomic changes in endothelial cells of Alveolus Chips at day 8 vs day 0 of culture. (**b**) Volcano plot of DEGs showing the transcriptomic changes in endothelial cells of Alveolus Chips at day 14 vs day 8 of culture. (**c**) Increased expression of several Wnt ligands in endothelial cells on-chip during Alveolus Chip maturation.

**Figure S5.**
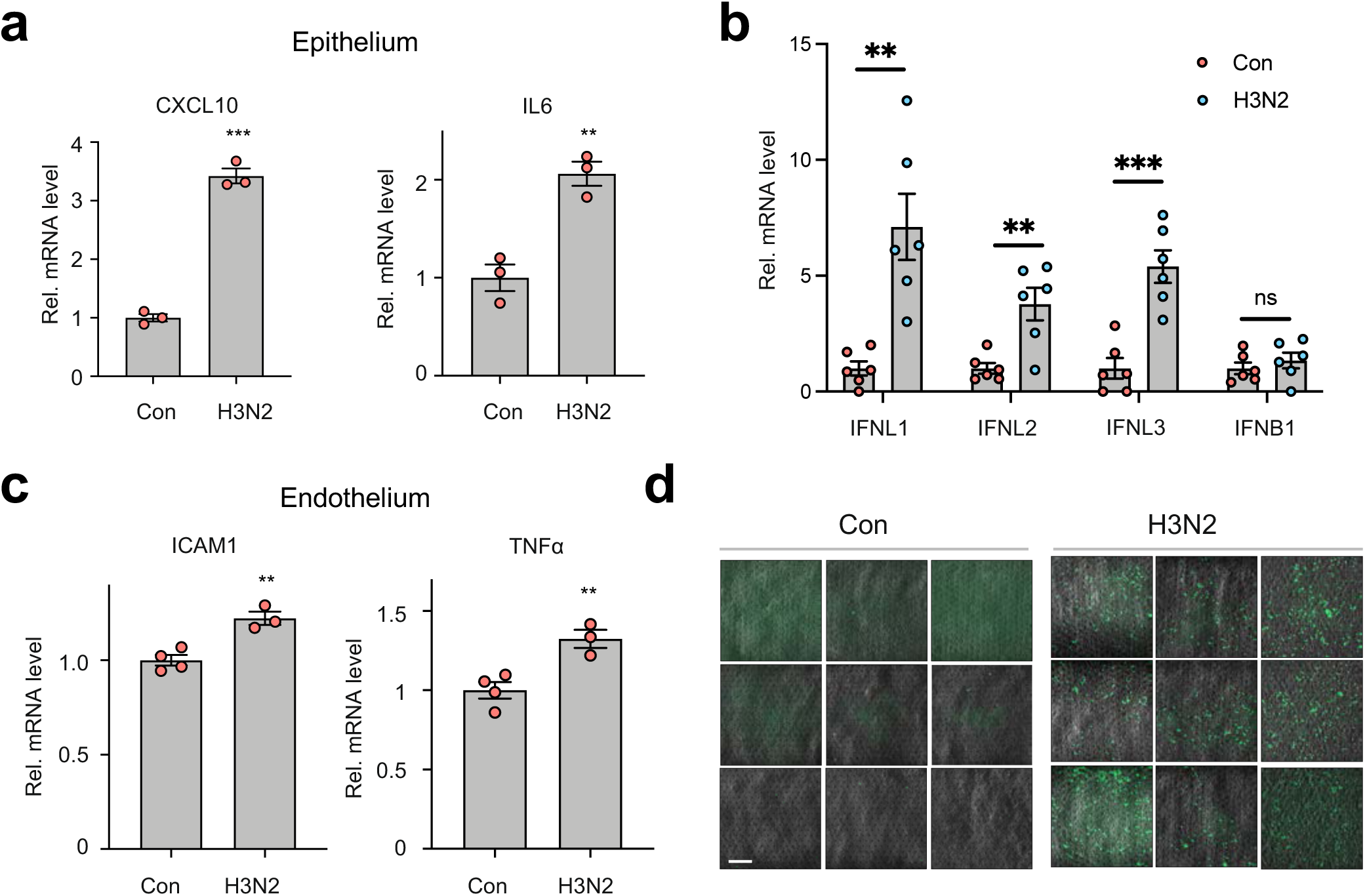
Host immune responses to influenza infection in Alveolus Chip. (**a**) Graphs showing relative mRNA levels of CXCL10 and IL-6 in epithelial cells of Alveolus Chips uninfected (Con) and infected with HK/68 (H3N2) (MOI = 1), measured by qPCR at 48 h post-infection. Data are shown as mean ± SD; n = 3 biological replicates; unpaired two-tailed t-test, **p<0.01, ***p<0.001. (**b**) Graph showing the relative mRNA levels of IFNL1, IFNL2, IFNL3, and IFNB1 in epithelial cells of Alveolus Chips uninfected (Con) and infected with HK/68 (H3N2) (MOI = 1), measured by qPCR at 48 h post-infection. Data are shown as mean ± SD; n = 6 biological replicates; unpaired two-tailed t-test, **p<0.01, ***p<0.001. (**c**) Graph showing the relative mRNA levels of ICAM1 and TNFα in endothelial cells of Alveolus Chips uninfected (Con) and infected with HK/68 (H3N2) (MOI = 1), measured by qPCR at 48 h post-infection. Data are shown as mean ± SD; n = 3-4 biological replicates; unpaired two-tailed t-test, **p<0.01. (**d**) Images showing the adhesion of CellTracker Green-labeled PBMCs to endothelium of Alveolus Chips uninfected (Con) and infected with HK/68 (H3N2). Nine representative images were taken for each condition. Scale bar: 100 µm.

**Figure S6.**
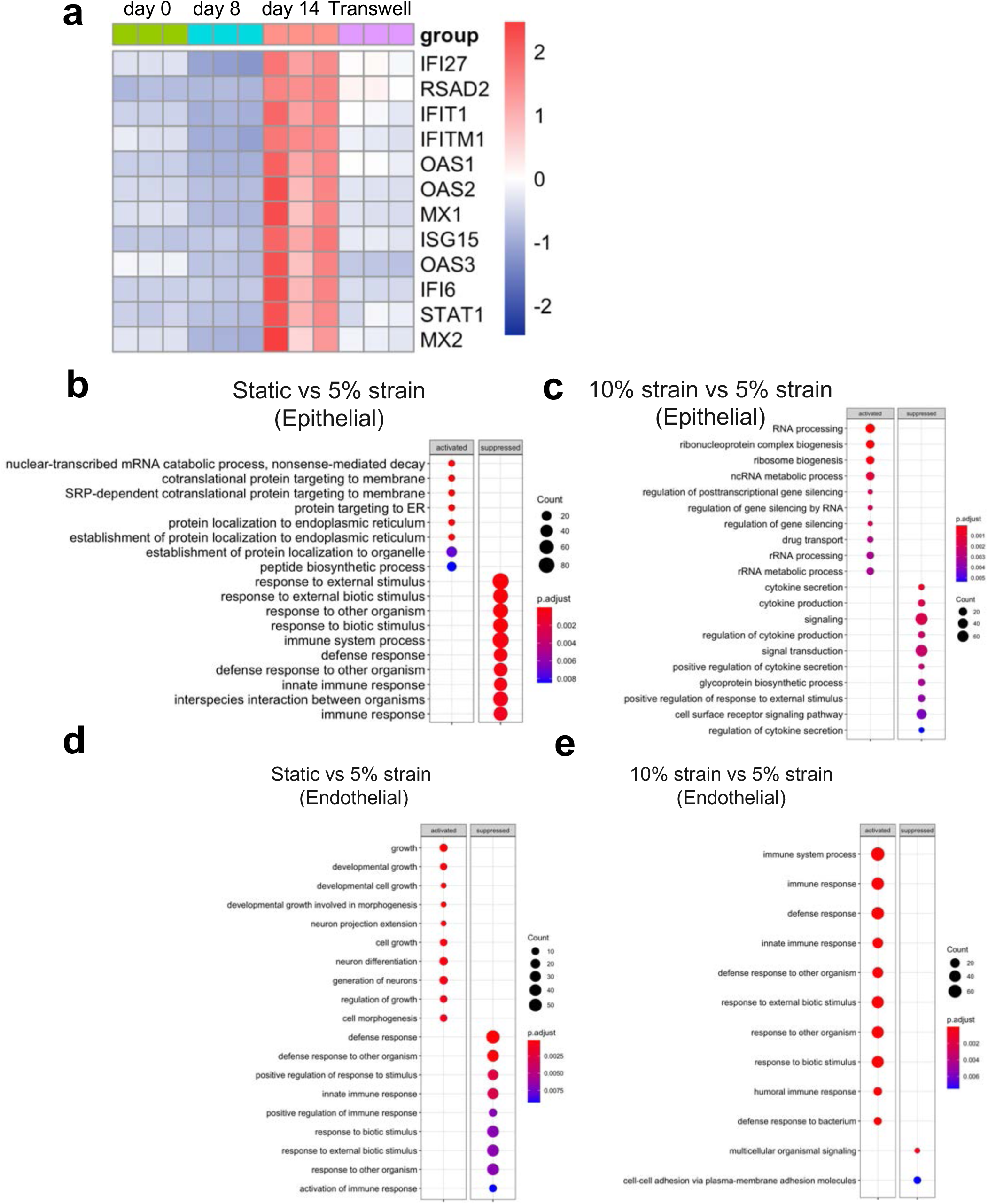
Cyclic mechanical strain activates the innate immune responses in Alveolus Chips. (**a**) Heat maps showing expression of interferon stimulated genes (ISGs) in epithelial cells of Alveolus Chips at different time points of culture or in transwell. (**b to e**) Gene ontology (GO) pathway-enrichment analysis of DEGs. **b, c, d, e** corresponds to Fig. 4 **c, d, f, and g**, respectively.

**Figure S7.**
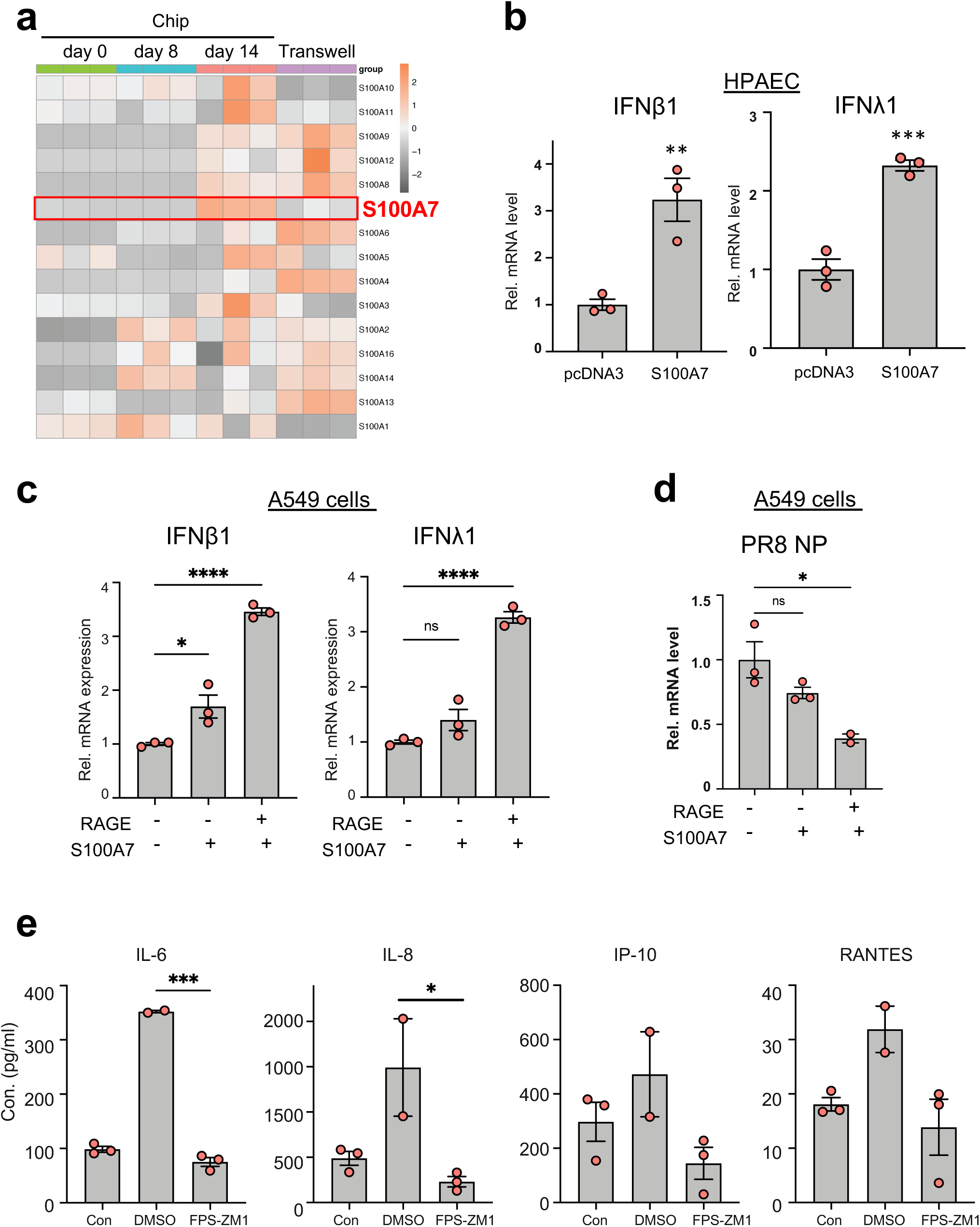
S100A7 induces innate immunity in Alveolus Chips. **(a)** Heatmap showing the specific increase of S100A7 among the S100 family genes during the Alveolus Chip differentiation. (**b**) Graph showing the mRNA levels of IFNβ1 and IFNλ1 in human primary alveolar epithelial cells (HPACE) at 48 hours after transfection with a plasmid expressing S100A7 or the pcDNA3 empty vector control. Data represent mean ± SD.; n = 3 biological replicates; unpaired two-tailed t-test, **p<0.01, ***p<0.001. (**c**) Graph showing the mRNA levels of IFNβ1 and IFNλ1 in A549 cells transfected with a plasmid expressing RAGE for 24 hours and then treated with culture supernatant containing S100A7 for 24 hours. Data are shown as mean ± SEM and statistical significance was assessed by one-way ANOVA with Tukey’s multiple comparisons test; ns, not significant; *p<0.05, ****p<0.0001. (**d**) Graph showing the levels of PR8 influenza virus NP mRNA in A549 cells transfected with a plasmid expressing RAGE for 24 hours, treated with culture supernatant containing S100A7 for 24 hours, and then infected with PR8 (MOI = 0.01) for another 24 hours. Data are shown as mean ± SEM and statistical significance was assessed by one-way ANOVA with Tukey’s multiple comparisons test; ns, not significant; *p<0.05. (**e**) Graphs showing the levels of cytokines in the vascular effluents of Alveolus Chips that were uninfected (Con), or infected with HK/68 (H3N2) (MOI = 1) in the presence or absence of 200 nM FPS-ZM1. Data are shown as mean ± SEM and statistical significance was assessed by one-way ANOVA with Tukey’s multiple comparisons test; n = 2-3 biological replicates; ns, not significant, *p<0.05, ***p<0.001.

